# Scx-positive tendon cells are required for correct muscle patterning

**DOI:** 10.1101/2021.01.03.424463

**Authors:** Yudai Ono, Tempei Sato, Chisa Shukunami, Hiroshi Asahara, Masafumi Inui

## Abstract

The elaborate movement of the vertebrate body is supported by the precise connection of muscle, tendon and bone. Each of the >600 distinct skeletal muscles in the human body has unique attachment sites; however, the mechanism through which muscles are reproducibly attached to designated partner tendons during embryonic development is incompletely understood. We herein show that Screlaxis-positive tendon cells have an essential role in correct muscle attachment in mouse embryos. Specific ablation of Screlaxis-positive cells resulted in dislocation of muscle attachment sites and abnormal muscle bundle morphology. Step-by-step observation of myogenic cell lineage revealed that post-fusion myofibers, but not migrating myoblasts, require tendon cells for their morphology. Furthermore, muscles could change their attachment site, even after the formation of the insertion. Our study demonstrated an essential role of tendon cells in the reproducibility and plasticity of skeletal muscle patterning, in turn revealing a novel tissue-tissue interaction in musculoskeletal morphogenesis.

**Graphical abstract:** 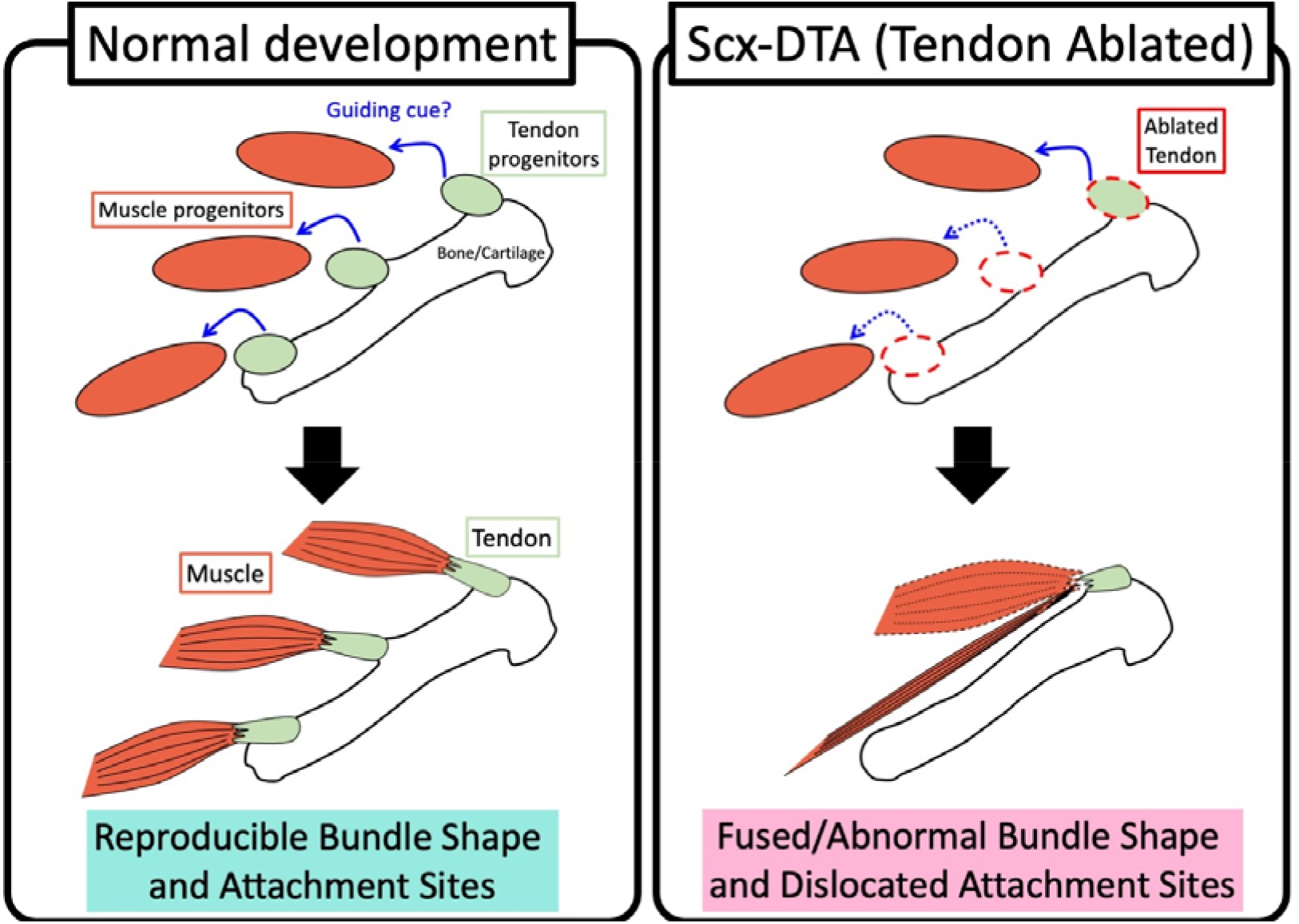

## Introduction

The musculoskeletal system is a complex multi-tissue system that consists of muscle, tendon and bone, as well as associating connective tissues. Elaborate body movements of vertebrate are supported by the precise shape, position and functional connections of these components. While the differentiation process of each component has been well studied (Asahara, Inui, & Lotz, 2017; Buckingham & Rigby, 2014; Kozhemyakina, Lassar, & Zelzer, 2015), their tissue integration process remain largely unexplored. Hundreds of skeletal muscles exist in the mammalian body; however, most are derived from the somites. Myogenic progenitor cells migrate long distances to destinations such as the limbs, form a precise bundle shape, and attach with appropriate tendons and bones during embryonic development (Buckingham et al., 2003; Comai & Tajbakhsh, 2014). It is well known that the migration of myogenic progenitor cells from somite to limb bud is guided by secreted cues such as HGF/SF or SDF-1 (Dietrich et al., 1999; Griffin, Apponi, Long, & Pavlath, 2010). However, mechanisms that regulate tissue integration between post-migration muscle cells and tendons and bones are not fully understood (Kardon, 2011; Schweitzer, Zelzer, & Volk, 2010). Considering the distinct cellular origin (i.e., muscles from the somites and tendons from the lateral plate) and large number of integrations to be formed in limited time and space, it is reasonable to assume that the local tissue-tissue interactions take place between myogenic progenitor cells and the surrounding cell/tissue. Indeed, several factors, including transcription factors, ECM, muscle connective tissue cells (MCTs) are reported to regulate the patterning of post-migrating myogenic cells in vertebrate limbs (Besse et al., 2020; Hasson et al., 2010; Helmbacher & Stricker, 2020; Kardon, Harfe, & Tabin, 2003; Kutchuk et al., 2015; Rodriguez-Guzman, Montero, Santesteban, Gañan, Macias, & Hurle, 2007b; Swinehart, Schlientz, Quintanilla, Mortlock, & Wellik, 2013). However, whether tendon cells, the intrinsic partner of skeletal muscles, have any role in instructing the muscle shape and patterning in mammals is unclear. To examine the role of tendon cells in the regulation of muscle patterning, we took a simple approach of linage-specific cell ablation and reduced the tendon cells in embryos. As a result, we found that muscle attachment patterns are significantly altered in the embryo with reduced tendon cells. Our results indicate that Scx-positive tendon cells have an important instructive role in spatially precise muscle attachment patterns, and are in turn essential for reproducible and robust musculoskeletal morphogenesis.

## Results and Discussion

### ScxCre mediated tendon ablation

To induce tendon cell-specific cell death, we mated a *ScxCre-L* Tg mouse (Sugimoto, Takimoto, Hiraki, & Shukunami, 2013b) with a *Rosa26-LSL-DTA* mouse (Voehringer, Liang, & Locksley, 2008) (Fig. 1A). As a result, we observed TUNEL-positive cells in tendon tissue of the tail and limb of Cre+/DTA+ embryo at E15.5, but not in control embryos (Fig. 1C, E arrowheads, Fig. S1 A-D). H&E staining of consecutive sections showed tissue ablation in the tail tendon tissue of Cre+/DTA+ embryo (Fig. 1B, D arrowheads). Cre and DTA dependent TUNEL-positive cells were observed as early as E12.5 in the forelimb, shoulder, intervertebral mesenchyme, or tail (Fig. S1 E-P). These results suggest that we could successfully induce cell death specifically in the tendon cells, from their early stage of differentiation. Hereafter, in this manuscript, we designate the mice with *ScxCre-L* Tg and *Rosa26-LSL-DTA* alleles as *“Scx-DTA”* mice.

**Figure 1.**
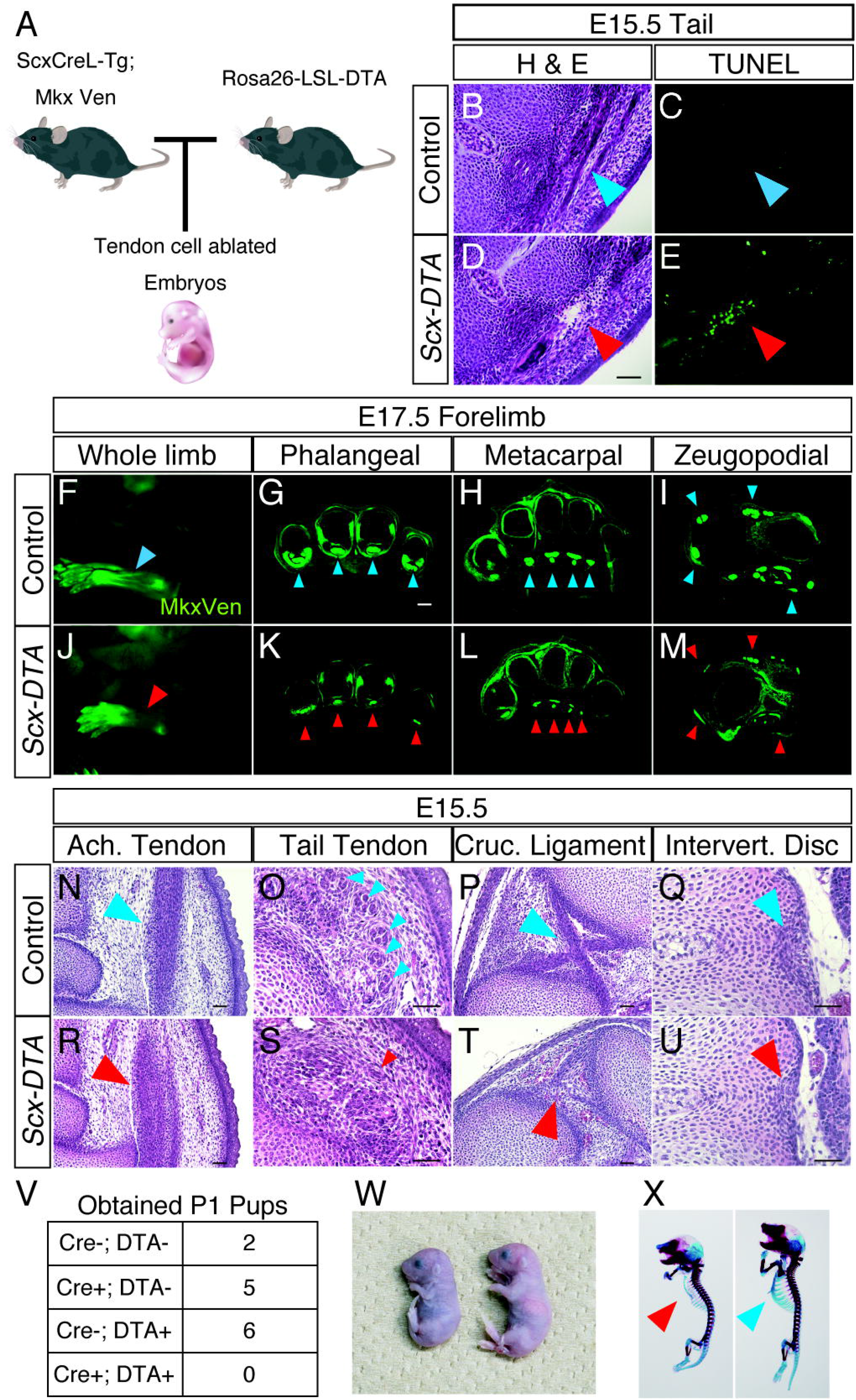
Tendon and ligament tissues are reduced in *Scx-DTA* mice. (A) A schematic drawing showing the generation of the *Scx-DTA* mouse. Illustrations are provided by ©2016 DBCLS TogoTV (B-E) H&E staining (B, D) and the TUNEL analysis (C, E) of the E15.5 embryonic tail. The blue arrowhead indicates the normal tail tendon and the red arrowhead indicates ablated tendon tissue. Scalebar: 100 μm. (F-M) The fluorescence signal from Mkx-Venus (MkxVen) knock-in reporter visualized the tendon tissue in E17.5 embryos. The blue arrowhead indicates normal limb tendons and the red arrowhead indicates ablated tendon tissue. Scalebar: 100 μm. (N-U) H&E staining of E15.5 embryonic tendon/ligament tissues. Ach. tendon, Achilles’ tendon; Cruc. Ligament, cruciate ligament; Intervert. Disc., intervertebral disc. Scalebar: 100 μm. (V) A table summarizing the genotypes of obtained P1 pups from crossing of *ScxCre-L* Tg and *Rosa26-LSL-DTA*. (W) The gross appearance of P0 pups. A *Scx-DTA* pup is shown on the left and a control pup is shown on the right. *Scx-DTA* pups were pale and the milk spot was not observed. (Y) Alcian-blue and alizarin-red staining showing the skeletal elements of P0 pups. An *Scx-DTA* pup is shown on the left and a control pup is shown on the right. The blue arrowhead indicates the normal ribcage and the red arrowhead indicates the reduced ribcage. For all the experiments, at least three embryos were examined (n>3) and consistent results were obtained. The figures show representative results.

Next, we examined the tendon tissue of *Scx-DTA* embryo with the *Mkx-Venus* knock-in allele (Ito et al., 2010). As shown in Fig. 1F and J, the long tendons in the zeugopod were reduced in E17.5 *Scx-DTA* embryo. The limb sections showed that the flexor digitorium profundus (FDP) tendon (Fig. 1G, K) and flexor digitorium sublimis (FDS) tendon (Fig. 1H, L) in the autopod, extensor carpi radialis tendons, extensor digitorium communis (EDC) tendons, or the palmaris longus tendon (Fig. 1I, M) in the zeugopod were greatly reduced in *Scx-DTA* embryos. The reduction of tendon tissue in other parts of the body, such as the Achilles tendon (Fig. 1N, R) or tail tendon (Fig. 1O, S) was also apparent. Furthermore, ligamentous tissue, such as the cruciate and patella ligaments were diminished (Fig. 1P, T) and the outer annulus fibrosus of the intervertebral disc was also reduced (Fig. 1Q, U). In sum, these results illustrated that our approach reduced the tendon and ligament tissues from the developing embryo, albeit not completely. The reductions of tendon tissue have been reported in mice with the knockout of tendon transcription factors (TFs), such as *Scleraxis* (*Scx*), *Mohawk* (*Mkx*) or *Egr1* (Guerquin et al., 2013; Ito et al., 2010; Murchison et al., 2007). The degree of tendon/ligament reduction in *Scx-DTA* mice was as severe as or even more severe than those observed in these TF mutants. Namely, the cell death was observed as early as E12.5 in *Scx-DTA* embryo, which is earlier than the reported tendon reduction in TF mutants. In addition, while the tendon reduction was seen mainly in the long tendons in TF mutants, most of tendon and ligament tissues were reduced in *Scx-DTA* embryos. The Incomplete loss of tendon tissue was probably due to the penetrance of Cre activity(Comai, Sambasivan, Gopalakrishnan, & Tajbakhsh, 2014), or continuous recruitment of *Scx*-positive tendon cells from limb mesenchyme population (Huang et al., 2019; Shwartz, Viukov, Krief, & Zelzer, 2016). As tendon cell death occurs in a period that overlaps the individuation of tendon from anlage (Huang et al., 2015), this could also have an impact on tendon patterning. Indeed, the numbers of FDS or EDC tendons were reduced and outline of each of the tendons were indistinct in the *Scx-DTA* embryo (Fig. 1H, I, L, M)

We found that *Scx-DTA* pups died soon after birth (Fig. 1V, W). Scx-DTA embryo showed a defect in diaphragm (described below), which could cause an insufficient respiratory function in newborn *Scx-DTA* pups. In addition, as *Scx* is expressed also in non-tendon/ligament tissues such as the patella, rib cage, and bone ridges (Blitz, Sharir, Akiyama, & Zelzer, 2013; Sugimoto, Takimoto, Akiyama, Kist, Scherer, et al., 2013a), these *Scx*-positive skeletal elements are also reduced or lost in *Scx-DTA* embryos (Fig. 1X, Fig. S2). The loss of rib cage could also cause respiratory problems. Thus, in the present study, we only focused on embryonic development. The effects of skeletal abnormality are also discussed (see below).

### Muscle patterning was altered in tendon-ablated embryos

Next, we examined whether the patterning (i.e., shape or attachment) of skeletal muscles was altered in *Scx-DTA* embryos. First, we applied whole-mount immunohistochemistry with a myosin heavy chain (MHC) antibody for the analysis of the forelimb of P0 pups. In control limbs, the muscles were located between their regular attachment sites (origin and insertion); for example, the deltoid muscle originates from spine of the scapula and inserts into deltoid tuberosity (Fig. 2B, C blue arrowhead). In contrast, muscles in *Scx-DTA* mice showed changes in their attachment sites; for example, the insertion site of the deltoid muscle changed to the shoulder joint (Fig. 2F, G red arrowhead). The shapes/attachments of muscles in the zeugopod also changed, for example the extensor carpi ulnaris (ECU) and extensor digiti quarti/quinti (EDQ) muscles were clearly separated in the control limb (Fig. 2D arrowheads), but they were fused into a single muscle bundle in the *Scx-DTA* limb (Fig. 2H arrowhead). The extensor pollicis muscles, which are normally covered by superficial muscles, became visible (Fig. 2D, H arrows). Of note, despite their morphological change, most of the muscles in the zeugopod attached with tendons at their distal end.

**Figure 2.**
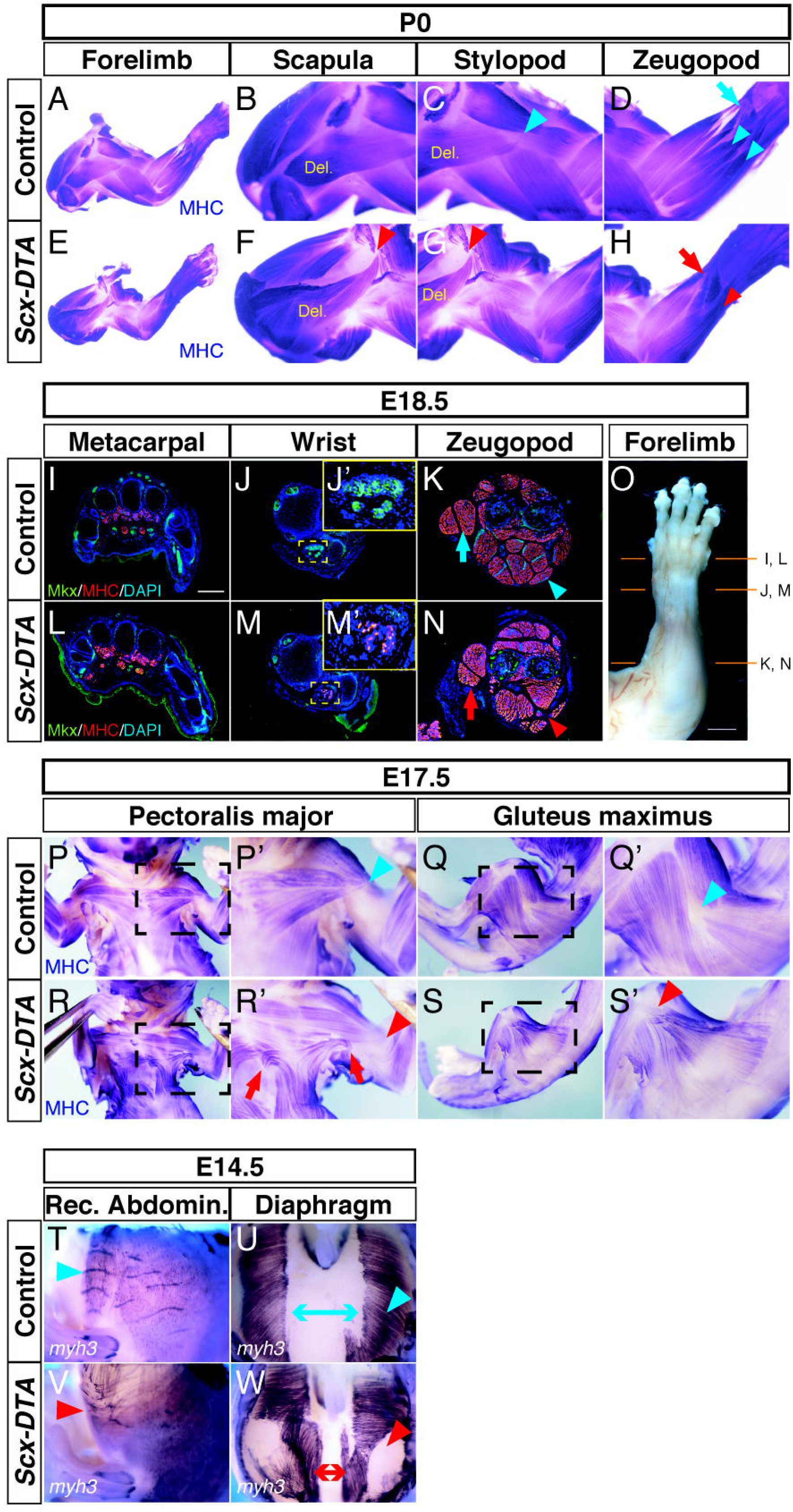
The muscle shape and position were altered in the *Scx-DTA* mouse. (A-H) Whole-mount immunohistochemistry of myosin heavy chain (MHC) in forelimbs of P0 pups. Del.: Deltoid muscle. The blue arrowhead in C indicates the normal insertion site of the deltoid muscle (deltoid tuberosity). The red arrowhead in G indicates an altered insertion site of the deltoid (shoulder joint). The arrows in D and H indicate the extensor pollicis muscles. The blue arrowheads in D indicate separated ECU and EDQ muscles. The red arrowhead in H indicates fused ECU/EDQ muscle. (I-O) Immunofluorescence of MHC and MkxVen (using α-EGFP) in the forelimbs of an E18.5 embryo. The blue arrowhead in J’ indicates FDS tendons. The red arrowhead in M’ indicates distally dislocated FDS muscle. The blue arrowhead in K indicates FDS muscles that are missing in the *Scx-DTA* mouse (red arrowhead in N). The blue arrow in K indicates normal extensor carpi muscles in a control embryo. The red arrow in N indicates rearranged extensor carpi muscles in an *Scx-DTA* embryo. The photograph in O shows the position of sections I-N. Scalebar: 100 μm. (P-S) Whole-mount immunohistochemistry of MHC in E17.5 embryos. The blue arrowhead in P’ indicates the normal insertion site of the pectoralis major muscle (deltoid tuberosity). The red arrowhead in R’ indicates a dislocated insertion site of the pectoralis major muscle (elbow joint). The blue arrowhead in Q’ indicates the normal insertion site of the gluteus maximus muscle (gluteus tuberosity). The red arrowhead in S’ indicates a dislocated insertion site of the gluteus major muscle (knee joint). (T-W) Whole-mount in situ hybridization of *myh3* in the rectus abdominus (T, V, Rec. Abdomin.) and diaphragm (U, W). The blue arrowheads in T and U indicate the normal organization of abdominal muscles. The red arrowheads in V and W indicate a disorganized pattern of abdominal muscles. The arrows in U and W indicate the width of the central tendons. For all of the experiments, at least three pups/embryos were examined (n>3) and consistent results were obtained. The figures show representative results.

In sections of E18.5 forelimbs, muscles in the metacarpal position were grossly normal (Fig. 2I, L, O). However, FDS muscles were found in the wrist position of the *Scx-DTA* limb, where muscle was not observed in control embryos (Fig. 2J, M, O). Conversely, FDS muscles were missing from their normal position in the zeugopod of *Scx-DTA* mice (Fig. 2K, N, O arrowheads). Interestingly, similar distal dislocation of FDS muscle has been reported in *Scx*-knockout mice (Huang et al., 2013), indicating that this change in muscle position is due to the reduction of FDS and FDP tendons. In addition, the arrangement of the extensor carpi (longus/Brevis) and FDP muscles was changed (Fig. 2K, N arrows). On the other hand, muscles in the hindlimb generally showed a normal attachment pattern (Fig. S3).

Altered attachment sites were also observed in other muscles, especially in muscles connecting to the body trunk and limbs. For example, the pectoral muscles originate from the sternum and insert into the deltoid tuberosity in control embryos (Fig. 2P, P’ arrowhead); however, the insertion sites of these muscles were distally dislocated toward the elbow joint in *Scx-DTA* mice (Fig. 2R, R’ arrowhead). In a severe case, the pectoral minor muscle separated into two bundles (Fig. 2R, R’ arrows). Similarly, in *Scx-DTA* mice, the insertion site of the gluteus maximus muscle, which normally originates from the ilium and inserts into the gluteus tuberosity (Fig. 2Q, Q’ arrowheads) changed distally, toward the knee joint (Fig. 2S, S’). The organization of muscles in the trunk, such as the rectus abdominis, was also changed in *Scx-DTA* mice (Fig. 2T, V). In the diaphragm, the width of the central tendon was greatly reduced (Fig. 2U, W arrows) and the orientation of associated muscle fibers was changed (Fig. 2U, W arrowheads). In sum, the organization of skeletal muscles of many body parts was changed in *Scx-DTA* mice, suggesting that tendon cells are necessary for skeletal muscle to form a precise shape and attachment in the correct position. The role of tendon tissue in muscle patterning has been shown indirectly in other vertebrate species; for example, the surgical removal of the tendon primordia resulted in ectopic extension of the muscle into the knee joint in chicken embryos (Kardon, 1998). *Scx*-knockout zebrafish embryos showed abnormal muscle patterning (Kague et al., 2019). In mammals, muscle-tendon interaction has been studied in the opposite direction and the requirement of muscle cells for tendon formation has been reported (Brent, Braun, & Tabin, 2005; Pryce et al., 2009). Our results represent the first evidence that directly points to the importance of tendon cells in muscle patterning in mammalian embryos. Muscle patterning in limbs also depends on LPM-derived mesenchyme cells or muscle connective tissue (MCT) cells that are closely associated with the myogenic lineage (Hasson et al., 2010; Kardon et al., 2003; Vallecillo-García et al., 2017). As such, we considered whether MCT is affected in *Scx-DTA* mice. As shown in Fig. S4, the expression levels of MCT marker genes were unaltered in *Scx-DTA* mice, while tendon marker genes were clearly reduced. Also, *Cre* expression in *ScxCre-L* Tg is restricted in tendon cells and not detected in MCTs (Sugimoto, Takimoto, Hiraki, & Shukunami, 2013b). We concluded that the phenotype seen in *Scx-DTA* mice is primarily due to the loss of tendon cells.

### Myofibers but not myoblasts require tendon cells for patterning

Next, to elucidate the stage at which tendon-muscle interaction takes place, we monitored the location and shape of the myogenic cell lineage in a step-by-step manner, along with their migration and differentiation. First, we found that *pax3*-positive myogenic progenitor cells were induced in the somite and migrated normally into the limb buds at E11.5 (Fig.3A, D arrowheads). *Myogenin*-positive myoblast positions were grossly normal at E12.5 or E13.5 (Fig.3B, C, E, F), suggesting that myoblast segregation occurred in similar manner in control and *Scx-DTA* mice. Furthermore, *myh3*-positive myofibers were observed in same position in *Scx-DTA* and control embryos at E12.5 (Fig. 3G, J), indicating that myofiber differentiation is not dependent on tendon cells. Of note, we found that the *myh3* signal became significantly high at the distal/proximal ends of myofibers from E13.5 (Fig. 3H, I arrowheads). By comparing the expression of *myh3* and a mature tendon marker, *tenomodulin* (*Tnmd*) (Shukunami, Oshima, & Hiraki, 2001; Shukunami et al., 2018), we noted that this high *myh3* expression marks the boundary of muscle and tendon (Fig. S3). We therefore interpreted the *myh3* high region as surrogate of the muscle-tendon boundary. The shape of *myh3*-positive muscle bundles clearly differed between *Scx-DTA* and control embryos at E13.5 (Fig.3H, K). While the EDC muscle in the control embryo formed a sharp boundary at its distal tip (Fig. 3H), the distal tip of EDC in *Scx-DTA* embryo remained loose shape without the clear boundary formation (Fig. 3K). Moreover, while the ECU and EDQ muscle bundles in control embryos were clearly separated and formed two distal boundaries (Fig. 3H blue arrowheads), those muscles did not separate or form normal distal boundaries in *Scx-DTA* embryos (Fig. 3K red arrowheads). The muscle patterning defects became more apparent in E14.5, where EDC/EDQ/ECU muscles could not be distinguished (Fig. 3I, L). Immunohistochemistry with MHC antibodies confirmed that the shapes of MHC-positive muscle bundles became loose in *Scx-DTA* embryos (Fig. 3M-P) and the insertion sites of several muscles, including the triceps (Fig. 3 M, O) and deltoid muscles (Fig. 3N, P), were dislocated. These results indicate that tendon cells are dispensable for the migration, initial segregation, or differentiation of myofibers, but are required for the formation of proper muscle bundle morphologies and the precise location of the attachment site. Previous studies showed that tendon anlages are induced in muscles-less limbs normally; however the maturation of tendons requires muscles (Brent et al., 2005; Huang et al., 2015; Pryce et al., 2009). Our results together with those studies, indicate that muscle and tendon anlage are induced independently, but their connection and maturation is mutually dependent. The patterning of limb muscle is regulated by surrounding tissues at various time point (Helmbacher & Stricker, 2020). While the loss of the *Tbx5* in *Prx1*-positive mesenchyme altered the patterning of tendon and muscle from E12.5 (Hasson et al., 2010), *Osr2Cre* dependent MCT ablation affected the patterning of *MfoD*-positive muscle clusters at E13.0 (Besse et al., 2020). Our results implied that the role of tendon cells, which is to finalize the muscle bundle position and attachment, is played later in comparison to other connective tissues. It is likely that multiple inputs from surrounding cells regulate limb muscle patterning sequentially and in an overlapping manner.

**Figure 3.**
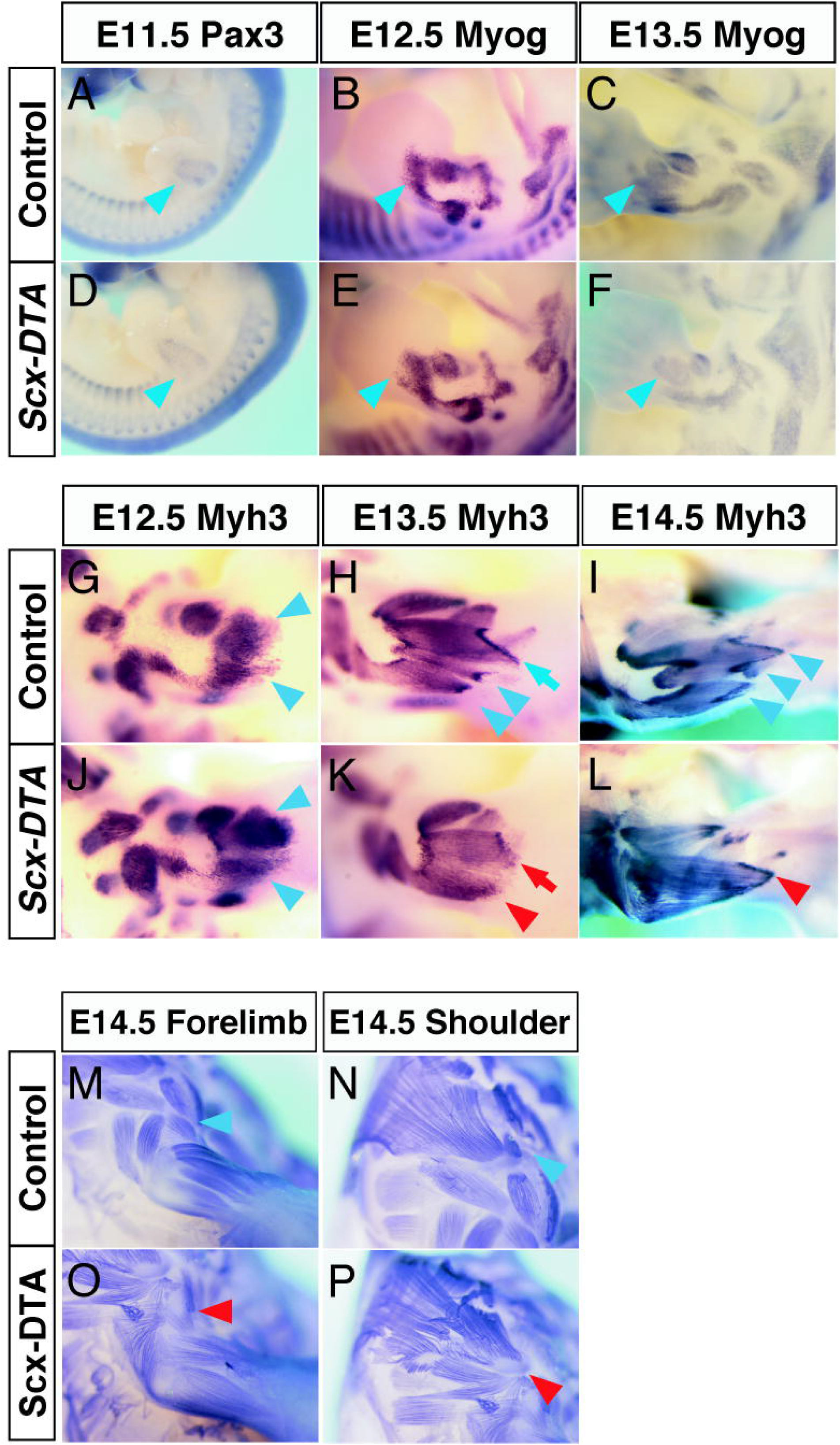
Loss of the tendon cells affects the patterning of myofibers. (A-F) A whole-mount in situ hybridization (WISH) analysis visualized the myoblast localization. The blue arrowheads in A and D indicate the normal migration of *Pax3*-positive myoblasts into the E11.5 limb. The blue arrowheads in B and E indicate the normal segregation of *Myog*-positive myoblasts in the E12.5 limb. The blue arrowheads in C and F indicate normal positioning of *Myog*-positive myoblasts in the E13.5 limb. (G-L) A WISH analysis visualizes myofiber patterning. The blue arrowheads in G and J indicate the normal location of *Myh3*-positive myofibers in the E12.5 limb. The blue arrowheads in H and I indicate the normal boundaries of the *Myh3*-positive FDC muscle bundle in E13.5 and E14.5 limbs. The red arrowheads in K and L indicate obscure or fused boundaries of the *Myh3*-positive FDC muscle bundle in E13.5 and E14.5 limbs. (M-P) Whole-mount immunohistochemistry visualized myosin heavy chain (MHC)-positive myofibers in E14.5 embryos. The blue arrowheads in M and N indicate the normal insertion site of the deltoid muscle (deltoid tuberosity). The red arrowheads in O and P indicate a dislocated insertion site of the deltoid muscle (shoulder joint). At least three embryos were examined in all the experiments (n>3) and consistent results were obtained. The figures show representative results

### Skeletal malformation or cell death are not the cause of muscle patterning defects

As *Scx* is also expressed in some skeletal elements, such as the rib and patella (Sugimoto, Takimoto, Hiraki, & Shukunami, 2013b), *Scx-DTA* mice showed several skeletal malformations, including loss of the rib cage or deltoid tuberosity (Fig. 1X, Fig. S2). To exclude the possibility that these skeletal malformation caused the muscle patterning defects, we examined the attachment pattern in embryos in which skeletal defects similar to those of *Scx-DTA* mice were reported (Braun, Rudnicki, Arnold, & Jaenisch, 1992; Kist et al., 2002). *Sox9* heterozygous (*Sox9^fl/-^*) embryos which lack the deltoid tuberosity were generated by mating *Sox9*^flox/flox^ and *Meox2-Cre* mice (Fig. S6B, D). Heterozygous *Myf5^Cre/+^* knock-in mice were intercrossed to generate homozygous *Myf5^Cre/Cre^* embryos, which resulted in rib cage malformation due to the loss of the Myf5 protein (Fig. S5F, H). In both cases the insertion sites of the pectoralis major muscles were not altered, despite the skeletal defects (Fig. S6 A, C, E, G). These results imply that the loss of skeletal elements is not the major cause of the muscle patterning defect seen in *Scx-DTA* mice. A previous study on the role of transcription factor *Lmx1b* also showed that the change of skeletal elements alone does not cause the muscle patterning change. Deletion of *Lmx1b* in a *Prx1*-positive lineage altered the dorso-ventral polarity of the whole limb, including the bone, tendon and muscle; however the loss of *Lmx1b* in the *Sox9*-positive lineage resulted in the change of bone, but not tendon or muscle polarity (Li, Qiu, Watson, Schweitzer, & Johnson, 2010).

Programmed cell death is one of the important driving forces of morphogenesis, such as digit formation or muscle belly segregation (Guha, Gomes, Kobayashi, Pestell, & Kessler, 2002; Rodriguez-Guzman, Montero, Santesteban, Gañan, Macias, & Hurle, 2007a). As such, the cell death induced in *Scx-DTA* mice could directly or indirectly affect the muscle attachment pattern. To explore this possibility, we examined the muscle patterning of the embryo where cell death is induced in muscle tissue by the *Myf5-Cre* dependent expression of DTA (*Myf5C^re/+^:Rosa26-LSL-DTA*, hereafter *“Myf5-DTA”*). In *Myf5-DTA* embryos, despite a severe reduction of muscle mass, the remaining pectoralis major muscle was correctly inserted into the deltoid tuberosity (Fig. S6I, J). This result indicated that excess cell death in the tissue does not necessarily cause muscle patterning defects.

### Dynamic change in muscle patterning after myofiber formation

The results shown above implied that the skeletal muscles define their attachment sites after myofiber formation (e.g., Fig. 3 G-L). We then asked if the position of muscle could be altered after the formation of insertion, which may provide skeletal muscle patterning further plasticity and robustness. To examine this point, we observed the position and insertion of gluteus maximus muscle at two time points E14.5 and E17.5. As shown in Fig. 4A and D, the distal tips of the gluteus maximus are located at the middle of the femur (i.e., the gluteus tuberosity) at E14.5 in both WT and *Scx-DTA* mice (Fig. 4A, D). A detailed section analysis confirmed that the gluteus maximus muscles formed insertions with gluteus tuberosity through tendon cells at this stage, although the tendon cells were reduced in *Scx-DTA* mice (Fig. 4B, C, E, F). As embryonic development proceeds, the junction of the gluteus maximus muscle/tendon matured and was firmly inserted into the gluteus tuberosity at E17.5 (Fig. 4G-I). On the other hand, the tip of gluteus maximus muscle of *Scx-DTA* mice was distally dislocated while most of the tendon and insertion into the gluteus tuberosity was lost by this stage (Fig. 4J-L). These results imply that the skeletal muscle is able to reposition its attachment site according to environmental changes, even after myofiber differentiation or the formation of initial attachment. Recent studies also reported the post-natal elongation or post-fusion repositioning of muscles (Gu et al., 2016; Huang et al., 2013), indicating that the plasticity of muscle positioning is higher than generally considered. We assume that this interaction between skeletal muscle and tendon provides musculoskeletal morphogenesis reproducibility and robustness to resist environmental or genetic perturbations.

**Figure 4.**
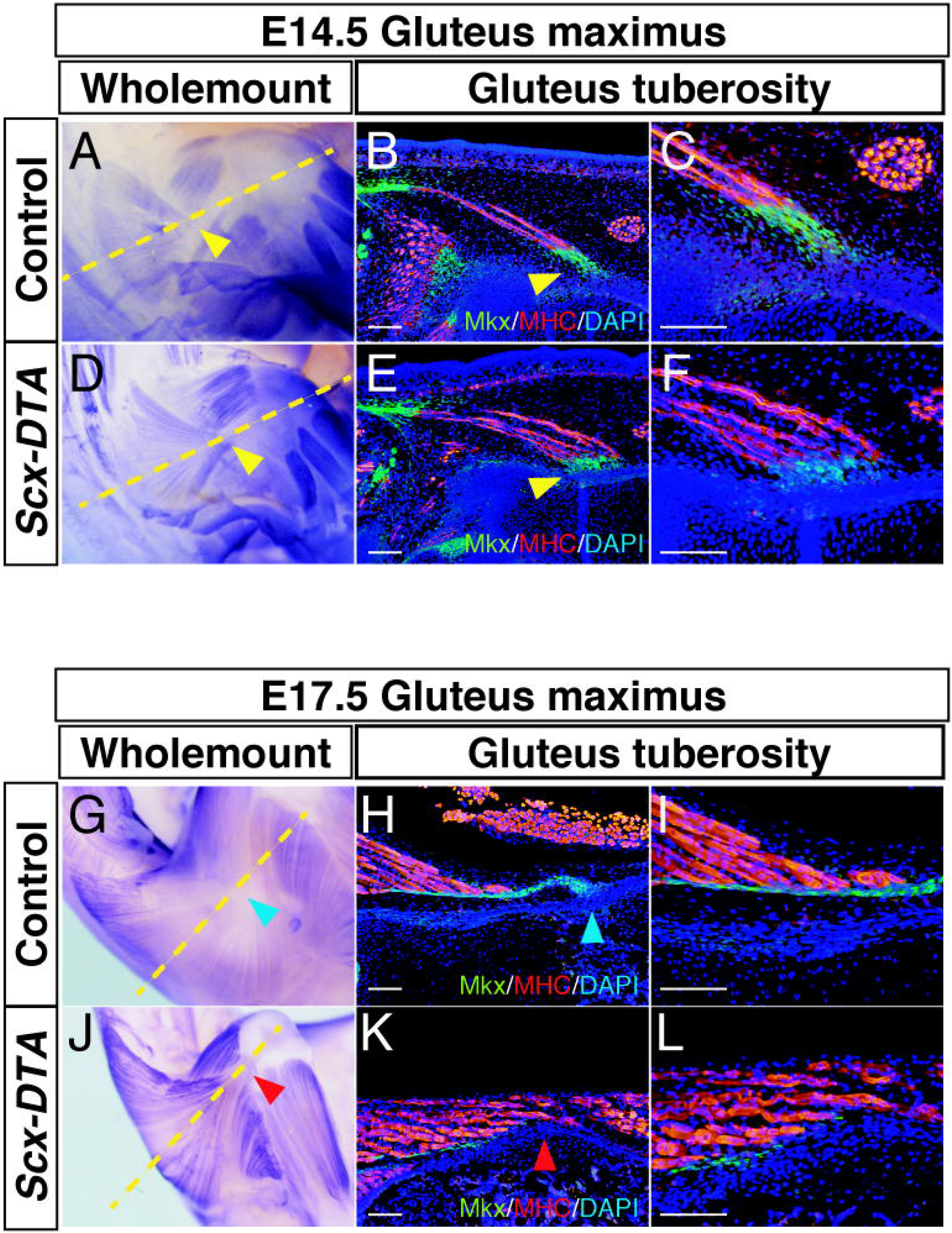
The attachment site of muscle was repositioned after myofiber differentiation. (A, D) Whole-mount immunohistochemistry visualized myosin heavy chain (MHC)-positive myofibers in the hind limb of E14.5 embryos. Dotted lines indicate the position of the section shown in B, C, E, and F. Arrowheads indicate the position of the gluteus tuberosities. (B, C, E, F) The immunofluorescence analysis of the E14.5 hindlimb. Arrowheads indicate the position of the gluteus tuberosities. Scalebar: 100 μm. (G, J) Whole-mount immunohistochemistry visualized myosin heavy chain (MHC)-positive myofibers in the hind limb of E17.5 embryos. Dotted lines indicate the position of the section shown in H, I, K, and L. The blue arrowhead in G indicates the normal insertion site of the gluteus maximus muscle (gluteus tuberosity). The red arrowhead in J indicates the dislocated insertion site of the gluteus maximus muscle (knee joint). (H, I, K, L) The immunofluorescence analysis of the E17.5 hindlimb. Arrowheads indicate the position of gluteus tuberosities. Scalebar: 100 μm. In all of the experiments, at least three embryos were examined (n>3) and consistent results were obtained. The figures show representative results

The molecular mechanism underlying this muscle-tendon interaction remains to be explored. In *Scx-DTA* mice, muscles in the zeugopod (i.e., the EDC/EDQ/ECU muscles) fused and attached to the remaining tendons at their distal end (Fig. 2H). The insertion sites of the muscles in the girdle (i.e., the pectoral and gluteus muscles) dislocated toward the joint regions, such as the shoulder, elbow or knee (Fig. 2, Fig. S7B). The remaining tendon cells were relatively abundant in the joint area of *Scx-DTA* embryos (Fig. 1F, J, Fig. S7A), probably due to the initial amount and continuous cell recruitment/differentiation from the *Scx*-negative cell population (Huang et al., 2019; Shwartz et al., 2016). The fact that muscle did not attach randomly to nearby bone but changed its morphology toward the distal tendon tissue implied a hypothesis that diffusible molecule(s) secreted from tendon cells could attract myofibers. Indeed, recent studies reported the involvement of retinoic acid in the formation of the extraocular functional unit (Comai et al., 2020), and FGF or BMP signaling were active at the interface of the embryonic tendon and muscle (Eloy-Trinquet, Wang, Edom-Vovard, & Duprez, 2009; Wang et al., 2010). However, cell adhesion molecule (Hasson et al., 2010) or ECM (Kutchuk et al., 2015) can also play parallel roles. Moreover, LPM-derived cells were recently suggested to convert their cell fate to the myogenic lineage at the tip of muscles and to regulate muscle patterning (de Lima et al., 2020). Clearly more studies are required to fully understand the molecular and cellular mechanisms of precise skeletal muscle patterning. We believe that revealing the molecular mechanism behind this process would shed light on broad areas of biology, such as the diversity of muscle patterning among species or regenerative medicine.

## Acknowledgements

We thank Drs. Scherer, Kist and Ohteki (TMDU) for providing the *Sox9 flox* and *Rosa26-LSL-DTA* mice. This Research is supported by AMED-CREST from AMED (JP20gm0810008), MEXT KAKENHI (19K06697), the Nakatomi Foundation, the Nakajima Foundation and the Takeda Science Foundation for M.I.

## Author Contributions

Y.O., T.S., C. S., and M.I. performed the experiments. H.A. and M.I. planned the study and wrote the manuscript.

## Declaration of Interests

The authors declare no conflicts of interest in association with the present study.

## Methods

### Mice

A *ScxCre-L* transgenic (Tg) mouse strain has been described previously (Sugimoto, Takimoto, Hiraki, & Shukunami, 2013b). A *Rosa26-LSL-DTA* mouse strain was kindly provide by Dr. Ohteki. The mouse was originally purchased from Jackson Laboratory (*B6.129P2-Gt(ROSA)26Sortm1(DTA)Lky/J*, strain#009669) by Dr. Ohteki and transferred under the permission of Jackson Lab. A *Mohawk-Venus* knock-in mouse was generated and described in our previous study (Ito et al., 2010). *Myf5Cre (B6.129S4-Myf5^tm3(cre)Sor^/J*, strain#007893) (Tallquist, Weismann, Hellström, & Soriano, 2000) and *Meox2Cre* mice (*B6.129S4-Meox2^tm1(cre)Sor^/J*, strain#003755) (Tallquist & Soriano, 2000) were purchased from Jackson Laboratory. A Sox9-flox mouse strain was kindly provided by Dr. Scherer and Dr. Kist (Kist, Schrewe, Balling, & Scherer, 2002). ICR mice were purchased from Sankyo lab-service (Tokyo, Japan). All animal experiments were approved by the animal experiment committees of Meiji University (approval No. IACUC17-0007) and the National Research Institute for Child Health and Development (approval No. A2004-003).

### Histological analyses

For paratlin sections, embryos were fixed with 4% paraformaldehyde (PFA) for 4°C overnight, dehydrated with methanol, cleared with xylene and embedded in paraffin For cryosection, embryos were fixed with 4% PFA for 4°C overnight, washed with PBS and embedded with OCT compound (Sakura Finetek, Osaka, Japan). The embryos were sectioned at 7 μm for hematoxylin and eosin (H&E) staining, and TUNEL and immunofluorescence analyses. The TUNEL analysis was performed using an In situ Death Detection Kit (Roche, Basel, Switzerland) according to the manufacturer’s instruction. The names of muscles and tendons appeared in sections were judged according to Watson et al. (Watson, Riordan, Pryce, & Schweitzer, 2009).

### Immunohistochemistry

The embryos were fixed with 4% PFA at 4°C overnight, dehydrated with methanol and rehydrated with PBS supplemented with 0.1% triton-X100 (PBSTx). The embryos were digested with 10 μg/ml Protease K at 37°C for 60 min, re-fixed with 4% PFA, blocked with 2% BSA/PBSTx and incubated with α-MHC antibody (Sigma-Aldrich, St. Louis, USA, My-32, 1:1000 in 2% BSA/PBSTx) at 4°C overnight. Then the embryos were washed 10 times with PBSTx, incubated with α-mouse IgG-AP conjugated (1:2000 in 2% BSA/PBSTx) (ab5880 Abcam, Cambridge, UK) at 4°C overnight, washed 10 times with PBSTx and the signal was developed in NBT/BCIP solution (Roche).

### Immunofluorescence

Paraffin or cryosections were prepared as described in the histology section, boiled in citric acid (pH 2.0) for antigen retrieval and stained with an α-MHC antibody (Sigma My-32, 1:1000) or α-GFP antibody (Abcam ab13970, 1:1000). α-mouse IgG-Cy3 (Jackson ImmunoResearch, West Grove, USA, 1:1000) or α-chicken IgG-Alexa488 (Thermo Fisher, Waltham, USA, 1:1000) were used as secondary antibodies.

### Whole mount *in situ* hybridization

Whole mount *in situ* hybridization (WISH) was performed according to the methods of Yokoyama et al. (Yokoyama et al., 2009). Briefly, the embryos were fixed with 4% PFA at 4°C overnight, dehydrated with methanol and rehydrated with PBS supplemented with 0.1% tween20 (PBST). The embryos were digested with 10 μg/ml Protease K at 37°C for 20-60 min (depending on the stage), re-fixed with 4% PFA, and hybridized with an anti-sense probe labeled with digoxigenin (DIG) or fluorescein in hybridization buffer at 65°C overnight. The embryos were washed with wash buffer at 65°C, blocked with 10% FBS/PBST for 2 hours, and incubated with α-DIG or α-fluorescein antibody conjugated with alkaline phosphatase (Roche, 1:2000 in 10% FBS/PBST) at 4°C overnight. Then the embryos were washed 10 times with PBST and the signal was developed in NBT/BCIP solution (Roche). For double *in situ* hybridization, DIG-labeled *tnmd* probe and fluorescein-labeled *myh3* probe were simultaneously hybridized. *Tnmd* was stained with α-DIG-AP antibody and NBT/BCIP substrate, post-fixed with 4% PFA, dehydrated with methanol, rehydrated with PBST and *myh3* was stained with α-fluorescein-AP antibody and INT (Roche)/BCIP (Roche) substrates. The sequences of the primers used for the amplification of probe are as follows: *myogenin* fw: 5’- ACCTGATGGAGCTGTATGAGACATC -3’, *myogenin* rev: 5’ -CATTTAGGTGACACTATAGCAGATGTGCACACTTGTCCAGG -3’, *myh3* fw: 5’ -CGTTTTGGACATTGCGGGTT -3’, *myh3* rev: 5’- ATGGACTCCCTCCTCTGCAT -3’.

### Skeletal preparation

The embryos were removed with their skin and internal organs, fixed with 100% ethanol, and serially stained with 0.03% alcian blue (Sigma-Aldrich) solution and 0.01% alizarin red (Sigma-Aldrich) solution. When embryos were used after whole-mount IHC, the embryos were post-fixed with 4% PFA, dehydrated with ethanol and stained as described above.

---

**Figure S1.**
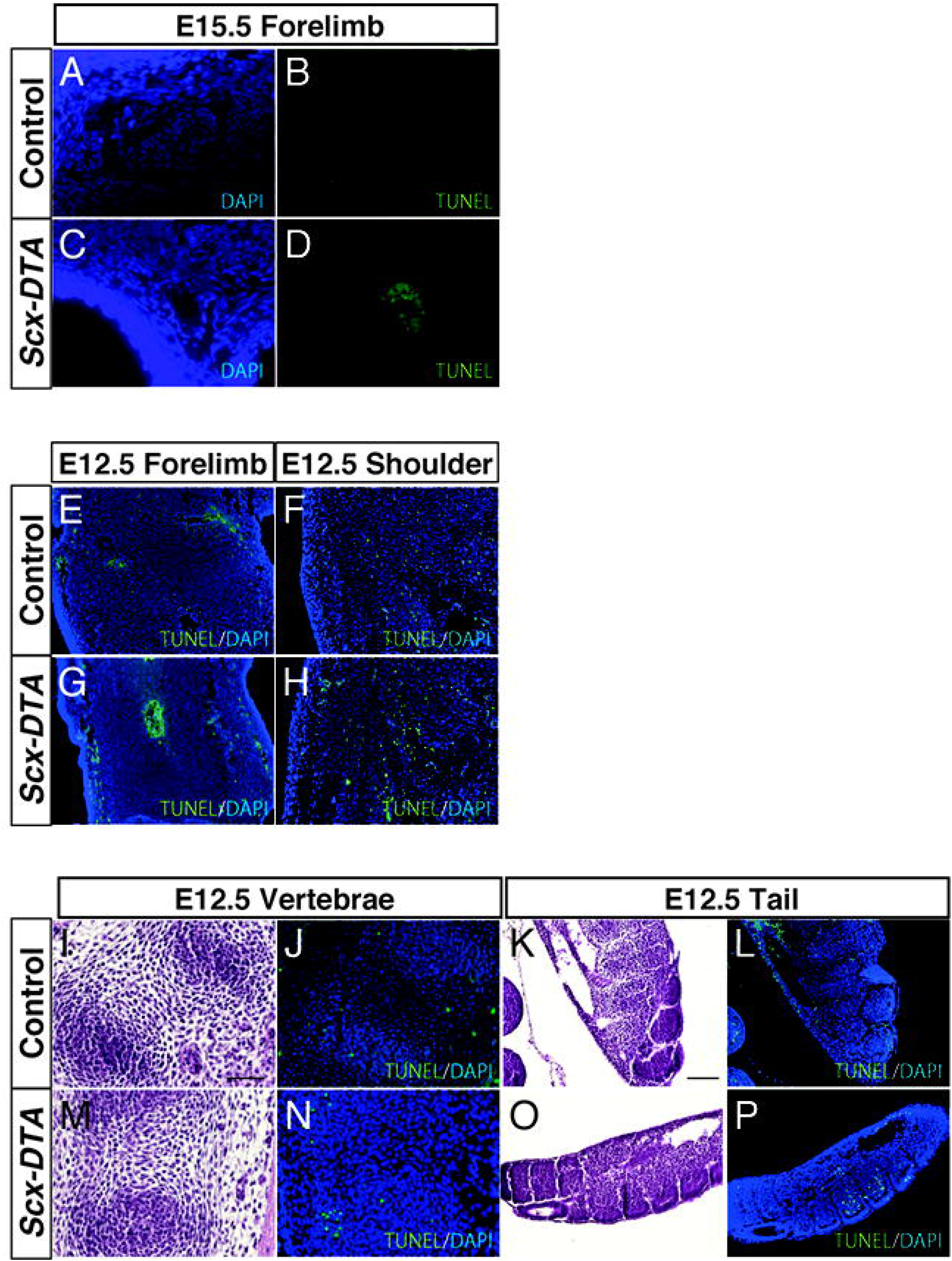
Tendon/ligament-specific cell death was observed in the *Scx-DTA* mouse. (A-D) A TUNEL analysis of the E15.5 forelimb. (E-H) A TUNEL analysis of the E12.5 forelimb and shoulder. (I-P) H&E staining and a TUNEL analysis of the E12.5 vertebrae and tail.

**Figure S2.**
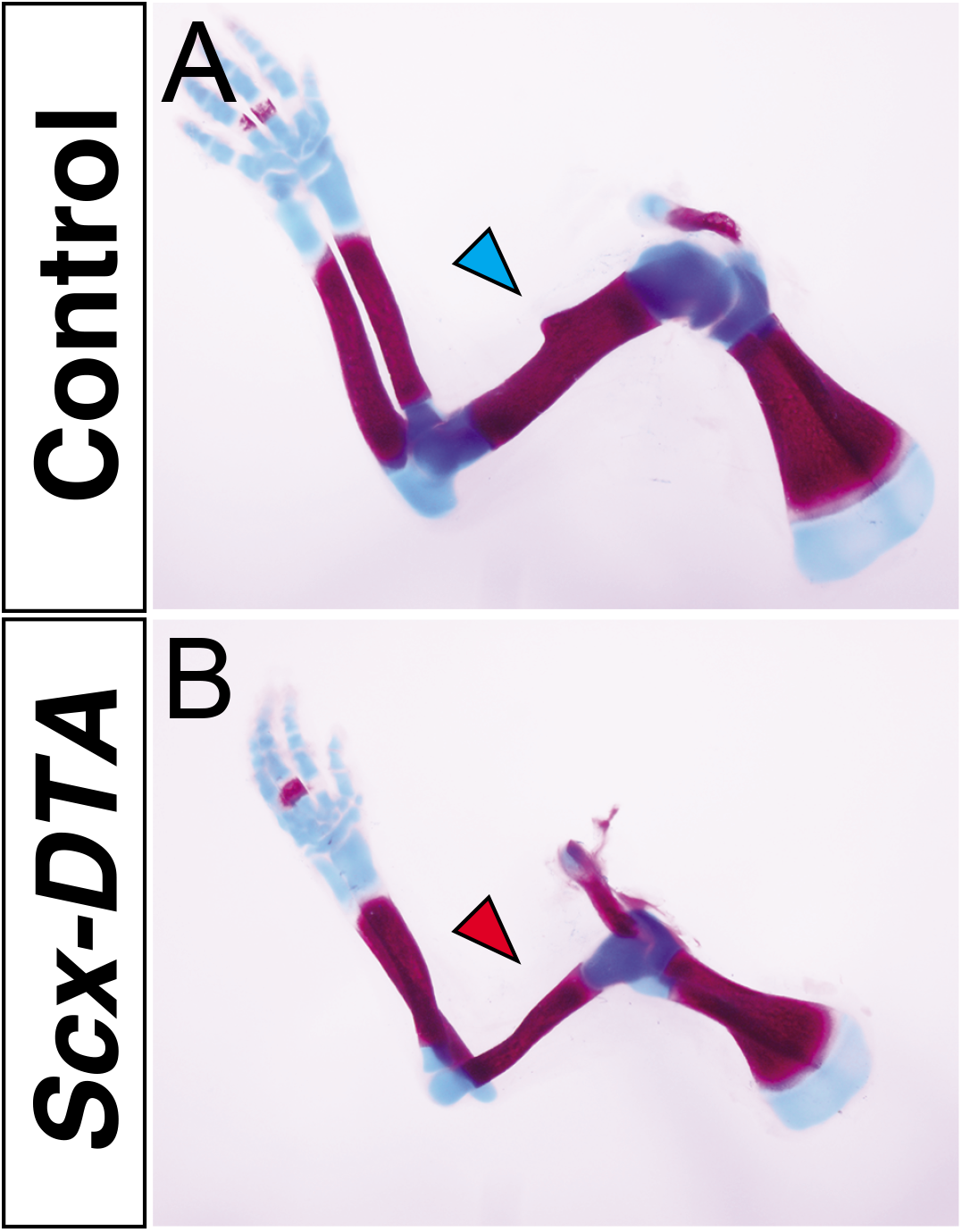
The deltoid tuberosity is missing in the *Scx-DTA* mouse. (A, B) Alcian-blue and alizarin-red staining of P0 pups. The blue arrowhead indicates the normal deltoid tuberosity in a control limb (A), which is missing in the *Scx-DTA* forelimb (B, red arrowhead)

**Figure S3.**
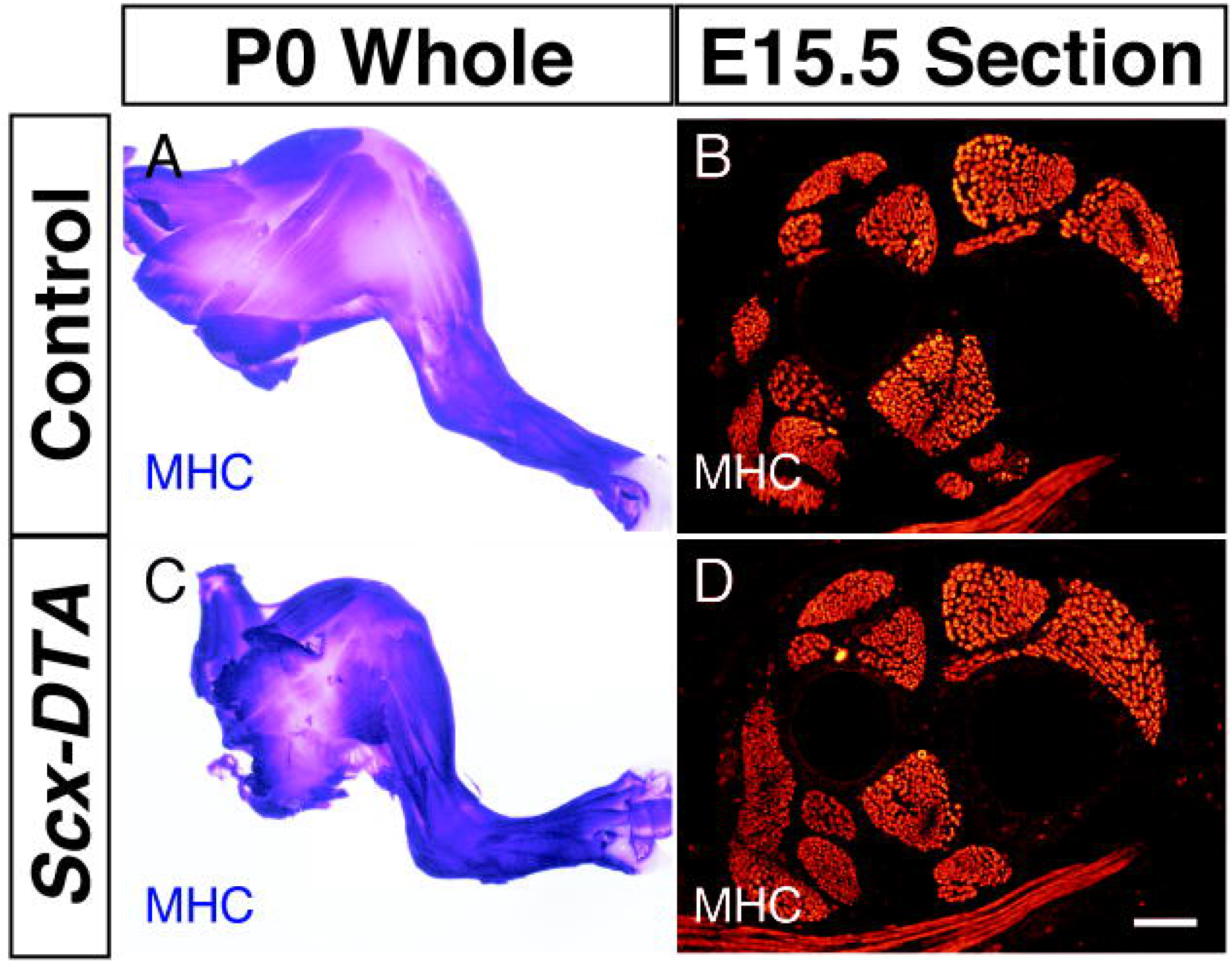
Muscle patterning was not affected in the hindlimb of the *Scx-DTA* mouse. (A, C) Whole-mount immunohistochemistry visualized MHC-positive myofibers in the P0 hindlimb. (B, D) An immunofluorescence analysis of MHC in an E15.5 hindlimb section. Scale bar: 100 μm

**Figure S4.**
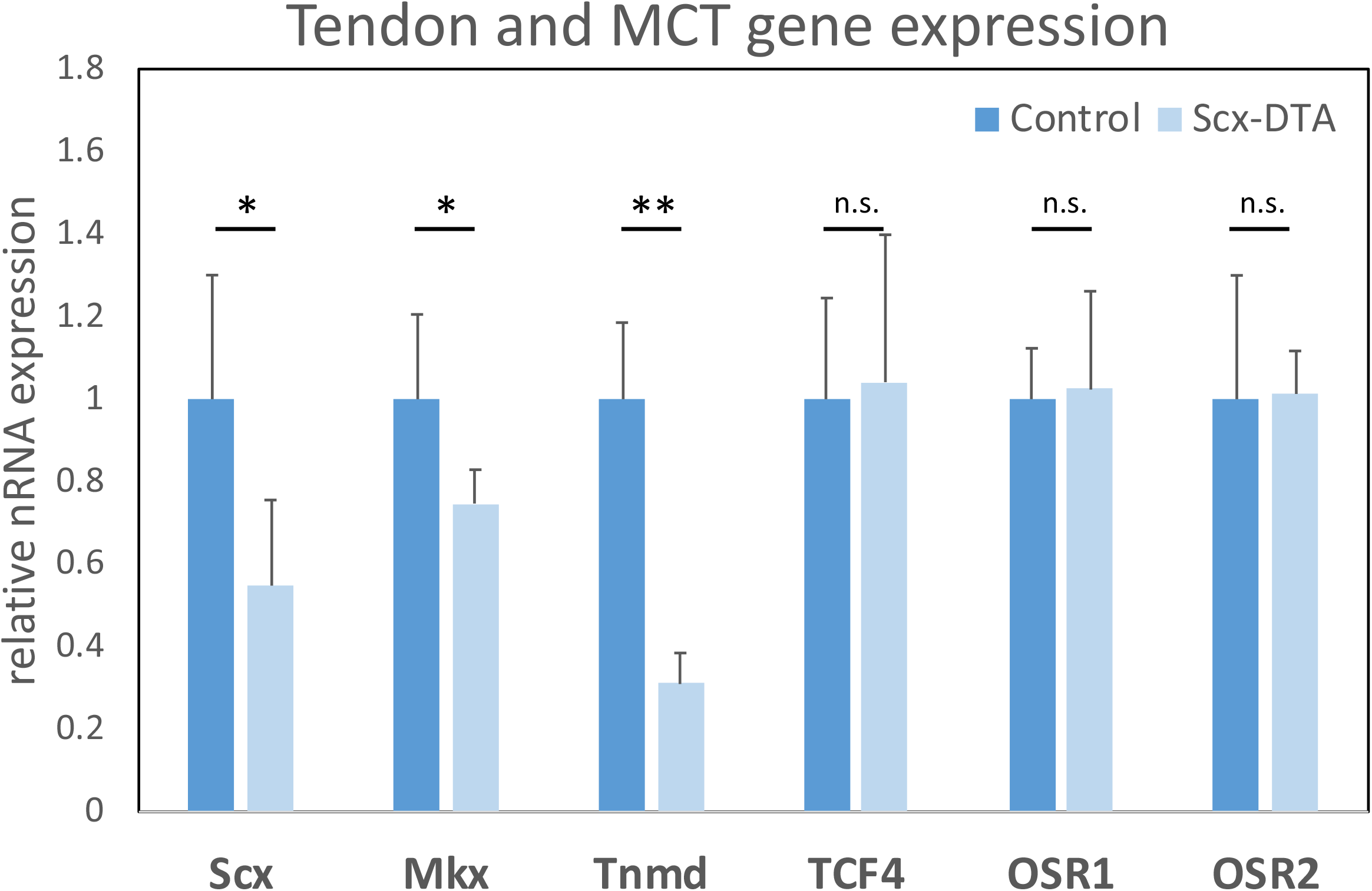
The MCT gene expression was not changed in the *Scx-DTA* mouse. Quantitative RT-PCR showed a significant reduction in the expression of tendon genes in the E14.5 *Scx-DTA* forelimb; however, the expression of MCT genes was not changed. Student’s t-test was performed. *p<0.05, **p<0.01, n.s.: no significant difference.

**Figure S5.**
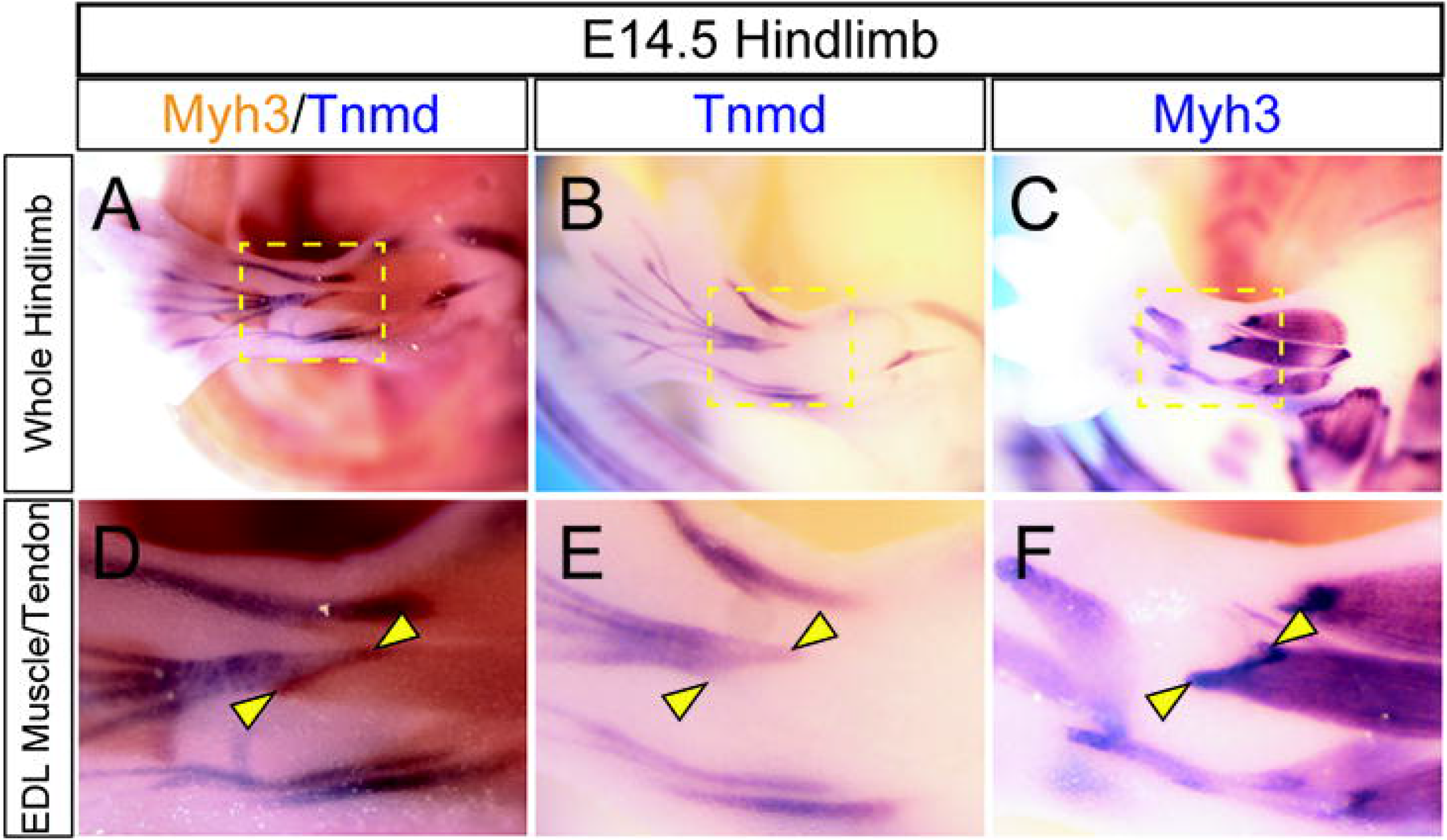
The high expression of *Myh3* marks the muscle-tendon junction. A whole-mount in situ hybridization (WISH) analysis visualized the *Myh3* and *Tnmd* expression in the E14.5 hindlimbs. In A and D, *Myh3* was stained with INT/BCIP (orange) and *Tnmd* was stained with NBT/BCIP (blue). Arrowheads indicate the putative junction between the EDL muscle and the EDL tendon.

**Figure S6.**
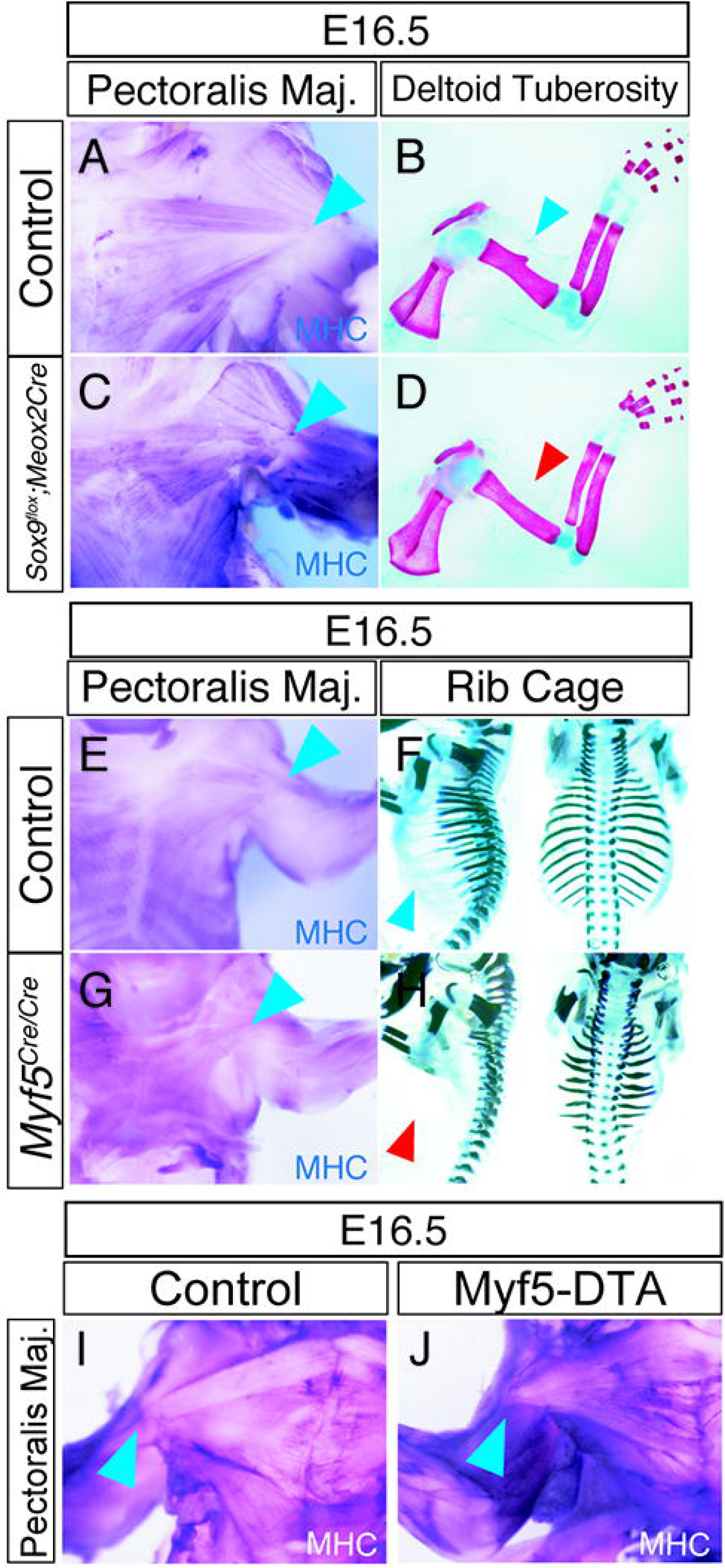
Skeletal malformation did not affect muscle patterning. (A, C) Whole-mount immunohistochemistry visualized MHC-positive myofibers in E16.5 pectoral muscles. The blue arrowheads indicate the normal insertion site of the pectoralis major muscles (deltoid tuberosity) in control and *Sox9* heterozygous embryos. (B, D) Alcian-blue and alizarin-red staining showing the skeletal elements of embryos shown in A and C. The blue arrowhead indicates the normal deltoid tuberosity in a control limb (B), which is missing in the *Sox9* heterozygous forelimb (D, red arrowhead). (E, G) Whole-mount immunohistochemistry visualized MHC-positive myofibers in the E16.5 pectoral muscles. The blue arrowheads indicate the normal insertion site of the pectoralis major muscles (deltoid tuberosity) in control and *Myf5^Cre/Cre^* embryos. (F, H) Alcian-blue and alizarin-red staining showing the skeletal elements of embryos shown in E and G. The blue arrowhead indicates a normal rib cage in a control embryo (E), which is diminished in *Myf5^Cre/Cre^* embryo (H, red arrowhead). (I, J) Whole-mount immunohistochemistry visualized MHC-positive myofibers in the E16.5 pectoral muscles. The blue arrowheads indicate the normal insertion site of the pectoralis major muscles (deltoid tuberosity) in control and *Myf5-DTA* embryos.

**Figure S7.**
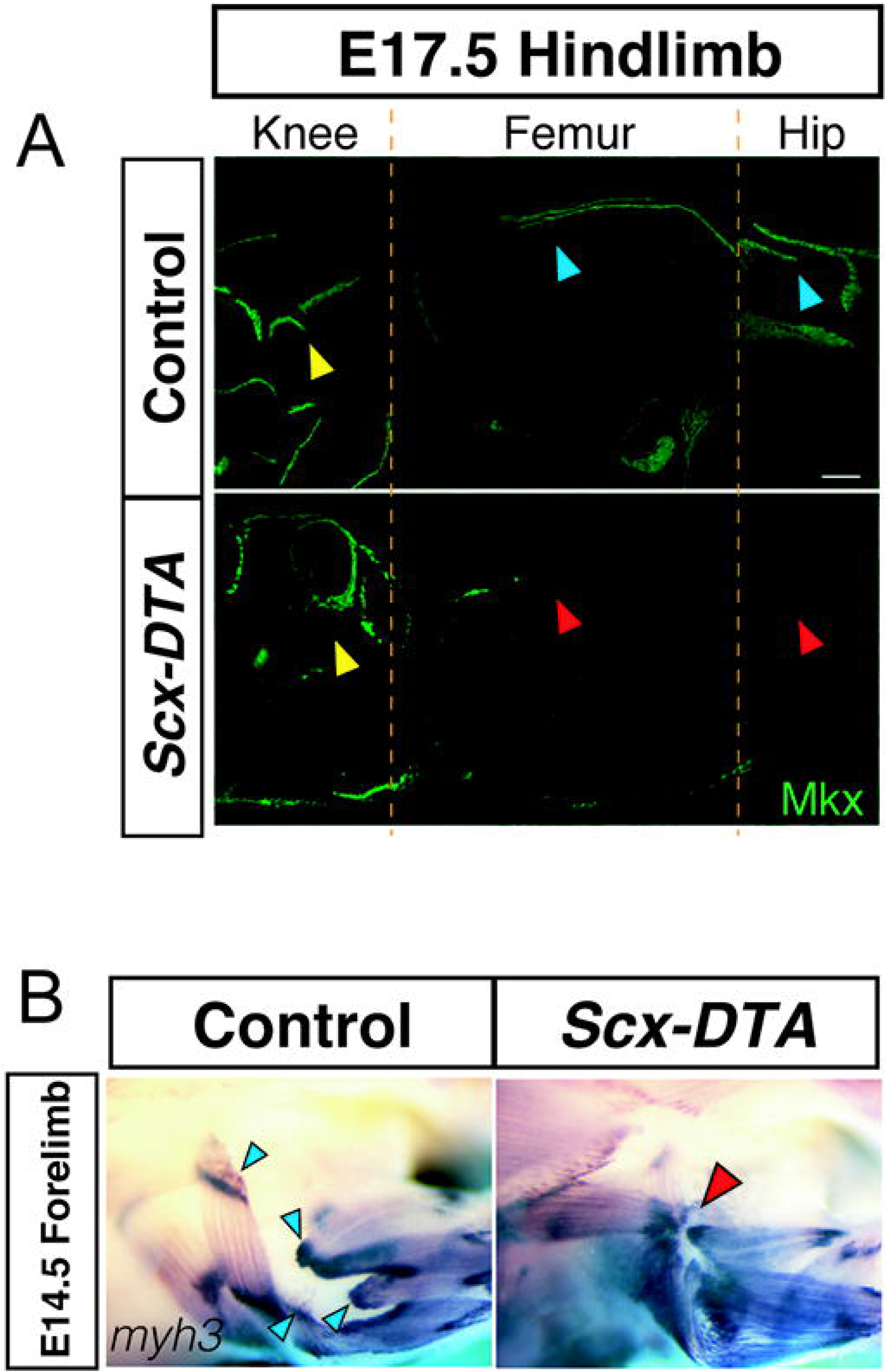
Dislocation of the insertion site toward the joint region where tendon cells are abundant. (A) The immunofluorescence analysis of MkxVen (using α-EGFP) in the E17.5 femur of control (upper panel) and *Scx-DTA* (lower panel) embryos. The blue arrowheads indicate the proximal and distal tendons for the gluteus major muscle in the control embryo, which are missing in the *Scx-DTA* embryo (red arrowhead). Yellow arrowheads indicate a comparable amount of tendon tissue in the knee joint of control and *Scx-DTA* embryos. (B) Whole-mount in situ hybridization of *myh3* in the E14.5 forelimb of control and *Scx-DTA* embryos. The blue arrowheads indicate the normal attachment sites of limb muscles. The red arrowhead indicates the accumulation of attachment sites to the shoulder joint.

## Notes

### Competing Interest Statement

The authors have declared no competing interest.

